# Association between recovering from tempo perturbations and reading measurements

**DOI:** 10.1101/2023.11.30.569275

**Authors:** Yi Wei, Roeland Hancock

## Abstract

A strong correlation between auditory temporal processing and reading proficiency has been consistently observed across clinical and nonclinical populations, spanning various age groups and languages. Specifically, rhythm sensitivity in both the music and speech domains has been considered to be fundamental for accurately tracking the hierarchical acoustic components in speech, playing a central role in the development of reading skills. However, the empirical validation of this hypothesis has primarily utilized stimuli with an isochronous underlying beat structure, which is limited in its ability to capture the nonlinearity inherent in the perception of timing within the speech and music domains. In our current study, we introduced perturbation stimuli and demonstrated a relationship between sensorimotor synchronization performance and reading measurements in the neurotypical adult population. Specifically, the current study highlighted that sensorimotor synchronization during the post-perturbation time window yields notably superior predictive value across a wide array of reading measurements when compared to the pre-perturbation time window, which, in contrast, did not predict reading measurements. Furthermore, our novel curve fitting analysis effectively captured the nonlinear aspects of participants’ sensorimotor synchronization performance when recovering from tempo perturbation, providing further insight into their auditory temporal processing abilities when responding to timing changes in auditory signals—a phenomenon commonly encountered in both speech and music contexts.

## Introduction

Despite the complex acoustic features of music and speech, the perception and cognition of these two domains come naturally and most of the time, seemingly effortlessly for humans. In contrast, reading is a skill that requires deliberate learning and often presents as a challenging and demanding task. The process of acquiring literacy is highly complex, involving the utilization of multiple cognitive abilities, which need to be supported by the establishment of efficient connections between various neural networks. Nonetheless, accumulating evidence from studies in music and speech rhythm perception consistently supports a strong association between auditory temporal processing in these domains and reading proficiency.

On the perception side, rhythm discrimination ability is positively associated with children’s reading skills (Anvari et al., 2002; Degé et al., 2015). Significant group differences in neural phase locking to stationary rhythmic stimuli have been found in children and adults with and without developmental dyslexia (Fiveash et al., 2020; Hämäläinen et al., 2012; Lizarazu et al., 2015; Soltész et al., 2013). However, Lizarazu et al. (2021) reported a failure to replicate some of these group differences.

On the production side, performance on sensorimotor synchronization to stationary stimuli has been positively linked to reading related measurements in English-speaking children (Woodruff Carr et al., 2014), French-speaking children (Lê et al., 2020), German preschoolers (Degé et al., 2015), and first grader Hungarian children (Kertész & Honbolygó, 2021). Similar relationships have been found in the English speaking adolescents (Tierney & Kraus, 2013b, 2013a), and both English and Mandarin speaking neurotypical adults (Wei et al., 2022). Longitudinal studies have provided additional evidence that preliterate temporal processing skills may be crucial in the development of phonological awareness and subsequent reading skills. These studies have shown consistent predictive power of music rhythm skills in later reading measurements, in English speaking children (David et al., 2007; Moritz et al., 2013), in Norwegian children (Lundetræ & Thomson, 2018), and French children (Dellatolas et al., 2009). Significant group differences in sensorimotor synchronization to metronome tasks have been found both in children (Bégel et al., 2022; Thomson & Goswami, 2008; Wolff, 2002) and adults (Thomson et al., 2006), with the exception of Rathcke & Lin (2021)’s study, in which they did not find an impairment in this task in adults with developmental dyslexia.

However, the causal link between auditory temporal processing and its influence on reading development remains a theoretical hypothesis. For example, Goswami (2011, 2015, 2019, 2022) proposed the Temporal Sampling Framework (TSF), suggesting that atypical neural phase entrainment to speech rhythm can affect children’s language development due to the inherited hierarchical timing structure in both neuronal activities and speech signals. Children and adults with developmental dyslexia have been the primary population targeted to test this relationship, given their characteristic clinical symptom of reading deficits. The experimental paradigms used to test this theory have mostly focused on establishing a relationship between perception and production of stationary stimuli and reading-related measurements (Fiveash et al., 2021; Ladányi et al., 2020). One critical link seemingly missing between this theoretical framework and experimental evidence is how we connect atypical behavioral/neural responses to stationary stimuli, such as a metronome, to atypical neural phase entrainment to speech rhythm.

Indeed, one often encountered argument is that the timing of musical notes is organized by an underlying framework of equal time intervals (Savage et al., 2015), while ordinary speech does not share this feature (Nolan & Jeon, 2014). However, we do not believe this argument is sufficient to invalidate the usage of musical stimuli. In fact, deviations from isochrony in musical rhythm are evident at both the level of individual notes, as observed in notes with fermatas or cadenzas, and at the level of meter structure, exemplified by intricate meters across various musical traditions (Polak et al., 2016). Neither notes nor meter in musical performance strictly adhere to periodicity (Palmer, 1997). Conversely, in the realm of perception, isochrony is discernible in speech rhythm during nursery rhymes and poetry recitation. Notably, in both domains, perceiving key event onsets does not invariably align with the acoustic properties of those events. In the domain of speech, listeners can perceive acoustically isochronous digit sequences as non-isochronous (Morton et al., 1976) and identify the P-centers of speech rhythm as distinct from both acoustic syllable onsets and intensity peaks (Marcus, 1981). Likewise, in the domain of music, individuals perceive a steady, isochronous pulse even when music produced by humans is inherently non-isochronous. Furthermore, they can perceive a pulse within certain types of syncopated rhythms where the pulse frequency is not physically reflected in the stimulus envelope (Chapin et al., 2010; Large et al., 2015; Tal et al., 2017).

This evidence suggests that the relationship between the physical acoustics in speech and music signals and our perception of rhythmicity in these two domains is not easily modelled by any linear system. We argue that to effectively assess the correlation between auditory temporal processing ability and reading proficiency, stimuli that are non-stationary yet capable of capturing the non-linear properties inherent in both music and speech perception and cognition should be employed. One example of a non-stationary stimulus suitable for sensorimotor synchronization tasks is a perturbation stimulus. A perturbation stimulus initiates with an entrainment time window, followed by an unexpected perturbation in period, phase, or both. When encountering the perturbation, participants adjust their synchronization to the tone onsets, resulting in a shift from a negative mean asynchrony (Schulze, 1992). Subsequently, participants recover from the perturbation, reverting to a negative mean asynchrony in their tapping behavior. Notably, the recovery time window exhibits an exponential decay due to the initial phase shift, enabling the capture of the nonlinear aspect of auditory temporal processing (Repp, 2002).

Not only can a perturbation paradigm capture nonlinearity in participants’ sensorimotor synchronization performance, but it can also provide additional insights into predictions from models such as TSF. For example, atypical rise time perception has been proposed to underpin phonological deficits observed in children with developmental dyslexia across languages. According to TSF, this atypical rise time perception results from atypical neural phase entrainment to slower time-scale rhythmic information in the speech signal. A perturbation paradigm can test not only the response to slower time-scale steady rhythm during the entrainment time window but also the response to timing changes that happen at a much faster time scale at the perturbation site. Specifically, we use exponential decay function to characterize participants’ responses to perturbation stimuli and model its association with their reading measurements. We argue that non-stationary stimuli, such as those of a perturbation paradigm, can better reflect the shared property of rhythm sensitivity in music and speech perception, therefore providing better theoretical and modeling power when testing the association with reading measurements.

## Methods

32 participants between 18-55 years old (M=25.53, SD = 9.98) were recruited from the University of Connecticut community. All participants reported being right-handed and native English speakers, with no history of neurological disorders. Some participants had experience playing instruments including string, keyboard, and wind instruments. Self-reported years of playing ranged from 0 to 16 years (M = 4.78, SD = 3.93), and years of music theory training ranged from 0 to 9 years (M = 1.22, SD = 2.39).

### Language Assessments

Participants completed tests of timed and untimed word reading, and timed passage reading fluency and comprehension. Timed word reading fluency was assessed using the Sight Word Efficiency and Phonemic Decoding Efficiency subtests from the Test of Word Reading Efficiency 2nd Ed (TOWRE-II, (Torgesen et al., 2012). Participants read a list of words or nonwords, respectively, out loud as fast and as accurately as they could within 45 seconds. Untimed word reading was assessed using the Word Attack and Letter-Word ID subtests from the Woodcock-Johnson III Tests of Achievement (Woodcock et al., 2001). In the Letter-Word ID and Word Attack subtests, participants read a list of words or nonwords, respectively, out loud, with no time constraints. Passage reading fluency and comprehension were assessed using the Gray Oral Reading Test (GORT, (Wiederholt & Bryant, 2012). The GORT is a timed task in which participants read aloud passages at different difficulty levels. Participants then answer comprehension questions. The GORT is scored on fluency and comprehension separately. The fluency score takes into account the time to finish reading the passage and the number of errors made during reading. The comprehension score is based on the number of correct answers on the comprehension questions. The raw scores of all the language assessments will go into regression models as dependent variables.

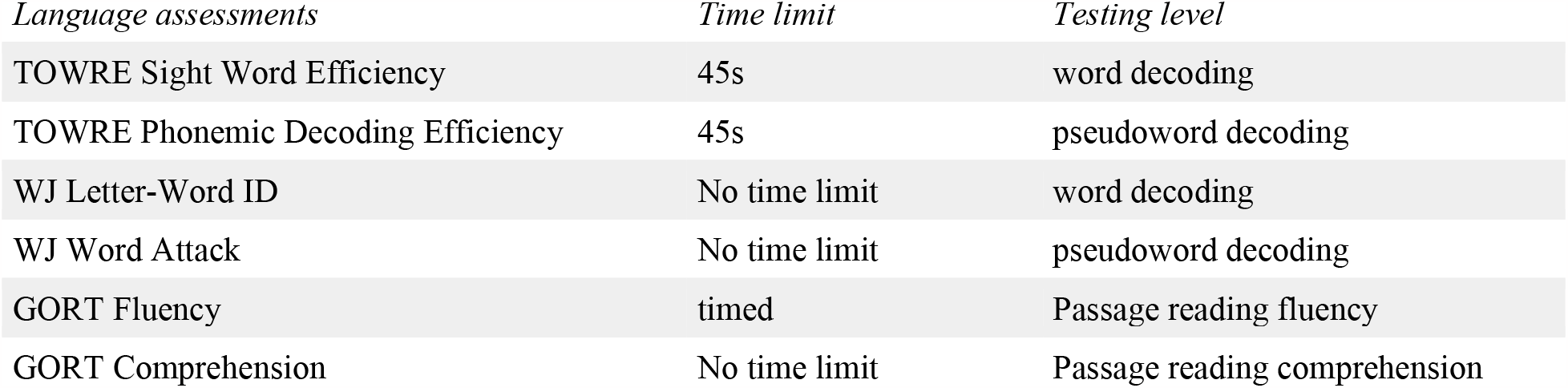

| *Language assessments* | *Time limit* | *Testing level* |
| --- | --- | --- |
| TOWRE Sight Word Efficiency | 45s | word decoding |
| TOWRE Phonemic Decoding Efficiency | 45s | pseudoword decoding |
| WJ Letter-Word ID | No time limit | word decoding |
| WJ Word Attack | No time limit | pseudoword decoding |
| GORT Fluency | timed | Passage reading fluency |
| GORT Comprehension | No time limit | Passage reading comprehension |

### Auditory stimuli

Isochronous rhythms with unpredictable tempo step perturbations were generated in MATLAB at 2Hz and 2.5Hz base tempi. All trials consisted of 20 tones. Each standard tone had an F0 of 200Hz with 5 harmonics. Tone length was 110ms, with 10ms ramp on, and 10ms ramp off time. A higher cue tone was 3/2 times the standard tone frequency, and a lower cue tone was 2/3 times the standard tone frequency. Inter-onset-intervals (IOIs) between the tones were perturbed pseudorandomly between the 9th and 12th tone, and all the IOIs were changed after the perturbation, creating a new tempo. The new tempi after the perturbation were changed by +20%, +25% (speeding up), -20%, and -25% (slowing down) of the base tempi. This created 8 rhythm conditions in total (2 tempi x 2 sizes x 2 directions) corresponding to 8 post perturbation tempi (1.5Hz, 1.6Hz, 1.875Hz, 2Hz, 2.4Hz, 2.5Hz, 3Hz, 3.125Hz). Each condition was repeated 48 times, creating 384 trials in total.

The participants’ synchronized finger-tapping responses were cued by the higher-pitched tone to start and a lower-pitched tone to end. The trials were divided into two categories: 1) early tapping trials, in which participants tap throughout the pre and post perturbation window. The start and end cue tones were randomly placed such that 6 tones before the perturbation site and 7 tones after the perturbation site were guaranteed to occur without a cue tone. Participants’ sensorimotor synchronization performance was analyzed with these trials. 2) late tapping trials, in which participants tap only during the post-perturbation window. The start cue tone occurred 9 tones after the perturbation site and tapping continued through the end of the trial. This approach allows the examination of neural entrainment to the auditory rhythm without requiring explicit sensorimotor synchronization. Early tapping trials and late tapping trials were interleaved in each of 16 blocks, with 24 trials in each block.

## Procedure

The whole experiment was completed in two separate sessions for each participant. In one of the sessions, participants’ anatomical T1-weighted structural MRI was collected on a Siemens Prisma 3 Tesla scanner. In the first session, participants completed the timed and untimed word reading fluency assessments. Participants then completed half (8 blocks) of the rhythm trials in a sound-attenuated booth. Stimuli were presented to participants through diotic ear inserts (Etymotic ER3) using Psychtoolbox 3 in MATLAB. EEG data was recorded at a sampling rate of 500 Hz with EGI’s Net Station Acquisition software, via a HydroCel 256-channel Geodesic Sensor Net and EGI NetAmps 400 amplifier. Participants were instructed to listen to the stimuli carefully and start tapping with their right index or middle finger as soon as they heard the high-pitched start cue tone. Participants were instructed to entrain their taps to every tone in the stimuli as accurately as they could until they either heard the low-pitched end cue tone, or the trial ended. Finger taps were recorded on a Roland percussion drum pad. Each trial was preceded by a two second pre-stimulus period of silent fixation on the screen. The whole entrainment session was recorded as one continuous EEG recording (∼ 45 mins). In the second session, participants first completed the timed passage reading fluency and comprehension assessments. Participants then completed the remaining 8 blocks of the rhythm task. The EEG results are outside the scope of this paper, and only the analysis and results of the behavioral portion are reported.

## Analysis

### Phase locking value analysis

To access phase stability of behavioral entrainment, for each participant, phase locking value (PLV) was calculated for pre-perturbation period and post-perturbation period separately, for each of the 8 conditions. PLV has been associated with reading measurements in neurotypical adults (Wei et al., 2022). To obtain PLV, first, the relative phase (in cycles) of each tap relative to the nearest stimulus tone (Large et al., 2002) was computed according to the formula:

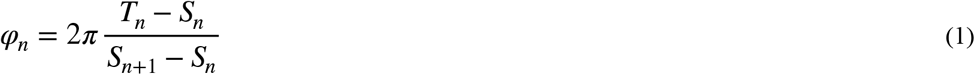

in which *T*_*n*_ is the time of tap, *S*_*n*_ is the time of the stimulus event closest to the tap *n. φ*_*n*_ is restricted to the range (-*π π*). PLV will then be computed according to the formula:

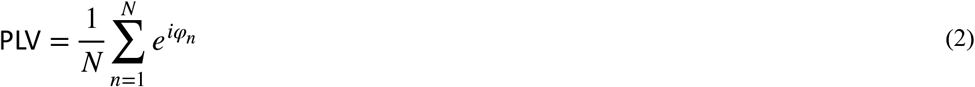

in which *N* = 8 is the number of taps during the post-perturbation period. The PLV values will be used in regression models to test their relationship with language measurements.

### Curve Fitting Analysis

To further investigate the dynamics of behavioral entrainment, a mono-exponential decay process of the relative phases over time after the perturbation was assumed, as it can capture three important parameters: the behavioral perturbation size (*θ*_0_), the recovery rate (*τ*), and the relaxation timing accuracy (*θ*_*a*_). Although more complex second-order systems have also been used to model post-perturbation phase (Large et al., 2002), the exponential model was selected based on initial visual inspection of the phase decay curves and desire to avoid possible overfitting.

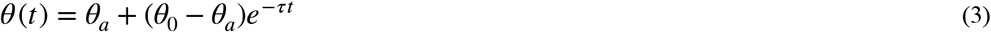

The MATLAB fitnlm function was used to fit the decay model (equation 3) to each trial using the relative phases of the 9 time points starting from the perturbation point. When perturbation direction is negative, the initial conditions of the model were set as *θ*_0_ = *π* /2, *θ*_*a*_ *=* − *π /4*, and. When perturbation direction is positive, the initial conditions were set as *θ*_0_ = − *π* /2, *θ*_*a*_ *=* − *π /4*, and.

### Regression analysis on the relationship between behavioral entrainment outcomes and reading measurements

For each of the behavioral outcomes (PLV and fitted decay parameters), a linear mixed effects regression model was first built to test its relationship with experimental manipulations (base tempi, perturbation size, and perturbation direction), after controlling for participants’ music background. For behavioral outcomes during the pre-perturbation (PLV-pre), only the manipulation “base tempi” was included as an experimental manipulation. In the case of *θ*_0_, only negative perturbation trials were included, since tapping at the perturbation site is only perturbed when perturbation direction is negative (speeding up). For behavioral outcomes during the post perturbation, which includes PLV-post, *θ*_*a*_, and *τ*, all three experimental manipulations were included.

Experimental manipulations that best account for the outcome variables were used to build a set of linear mixed effect models (full models) to test the relationship between each behavioral outcome and language measurements using the lmer function from LME4 R package (Bates et al., 2015). For the set of linear mixed effect models, behavioural outcomes (PLV-pre, PLV-post, *θ*_0_, *τ*, and *θ*_*a*_) were predicted by participants’ music background, identified significant experimental manipulations, language measurements, and the interaction between identified significant experimental manipulations and language measurements.

After the full model was built, stepwise regression analysis was applied to the full model to identify significant predictors that would enter the final linear mixed effect model. ANOVA type 3 analysis was then applied to the model found by the stepwise regression method, to determine significant relationships between behavioral outcomes, experimental manipulations, and language measurements.

## Results

### Behavioral entrainment analysis

Figure 1 shows average relative phase for each participant (thin lines) and for all participants (thick lines). Relative phase reaches a steady state before the perturbation (negative phase means participants tapped before the tone on average). At the site of the perturbation, relative phase is positively (fig1 a) or negatively displaced (fig 1c) (depending on perturbation direction). During the relaxation time window, relative phase relaxes back to the baseline, reaching steady state by the post-perturbation time window (Figure 1 shows only the largest perturbation, 25%). These results are consistent with previous experiments using this methodology (e.g., Large et al., 2002; Palmer et al., 2014; Rouse et al., 2016). Fig 1b and 1d show two representative participant’s curve fitting.

**Fig. 1.**
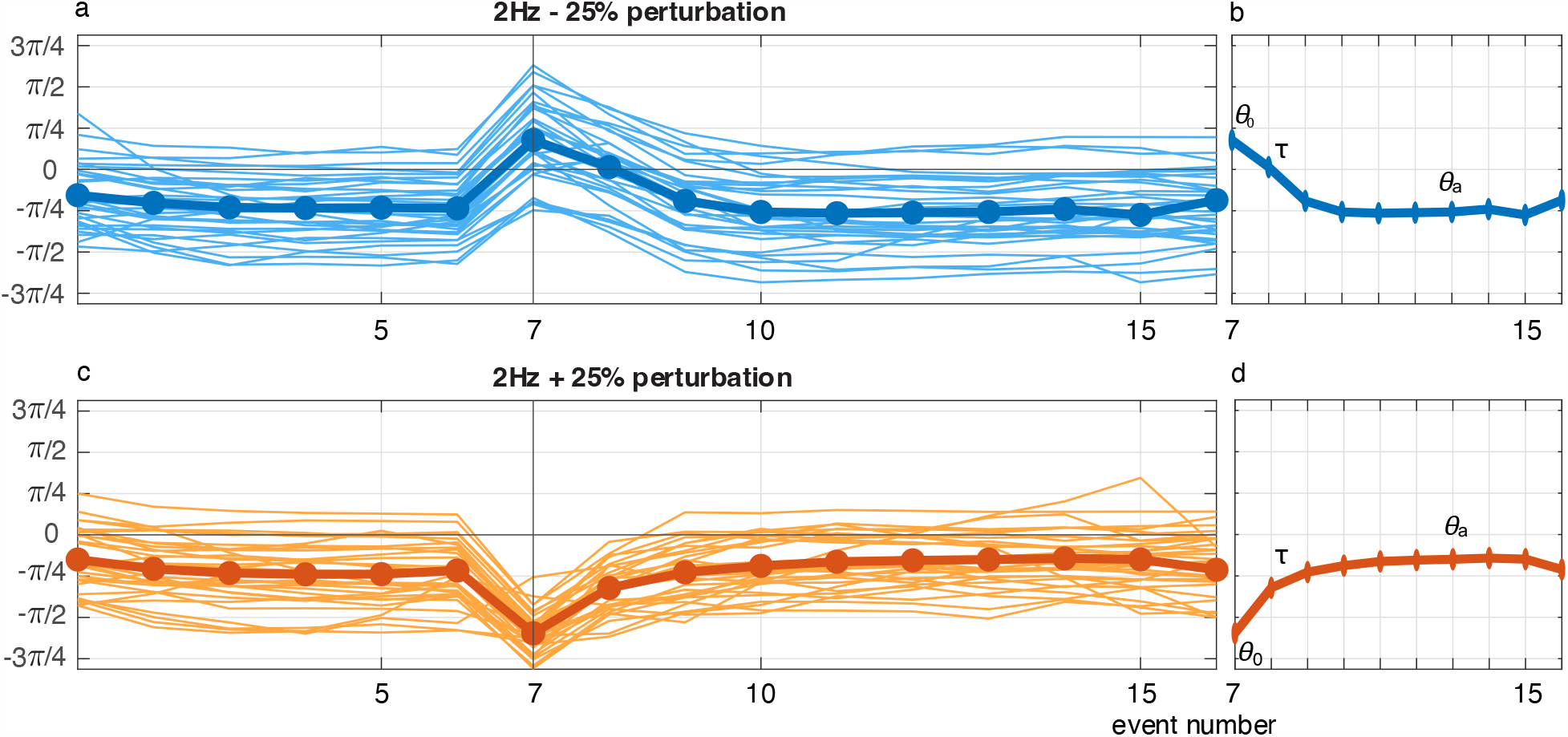
Behavioral entrainment in 2Hz (base tempi) 25% perturbation condition (a, c), and a representative participant’s curve fitting (b, d): *θ*_0_ indicates the size of perturbation in relative phase at the perturbation site, is the exponential decay rate, and *θ*_*a*_ indicates the asymptote of relative phase after recovery from perturbation.

### Relationship between experimental manipulation and behavioral entrainment outcomes

Models that tested the relationship between behavioral outcomes and experimental manipulation were as follows:

~~~
model <- lmer(PLV-pre ∼ age + instrument + theory + base tempi + (1|sub), data = data)
model <- lmer(*θ*_0_ ∼ age + instrument + theory + base tempi + perturbation size + (1|sub), data = data)
model <- lmer(PLV-post / *τ* / *θ*_*a*_ ∼ age + instrument + theory + base tempi + perturbation size + perturbation direction + (1|sub), data = data)
~~~

According to the model comparison, base tempi significantly predicted PLV-pre (*b* = -0.021, *sE* = 0.006, t = -3.61, p < 0.001). All three predictors (base tempi, perturbation direction, and perturbation size) significantly predicted PLV-post (base tempi: *b* = -0.026, *SE* = 0.010, t = -2.682, p < 0.01; perturbation direction: *b* = -0.017, *SE* = 0.005, t = -3.582, p < 0.001; perturbation size: *b* = -0.008, *SE* = 0.001, t = -8.778, p < 0.001). Base tempi and perturbation size significantly predicted *θ*_0_ (base tempi: *b* = 0.310, *SE* = 0.064, t = 4.862, p < 0.001; perturbation size: *b* = 0.050, *SE* = 0.006, t = 7.912, p < 0.001). Perturbation direction significantly predicted (*b* = -0.529, *SE* = 0.064, t = -8.315, p < 0.001), and *θ*_*a*_ (*b* = -0.348, *SE* = 0.040, t = -8.768, p < 0.001). Table 1 shows the relationship between experimental manipulations and behavioral measurements model outcomes.

**Table 1.**
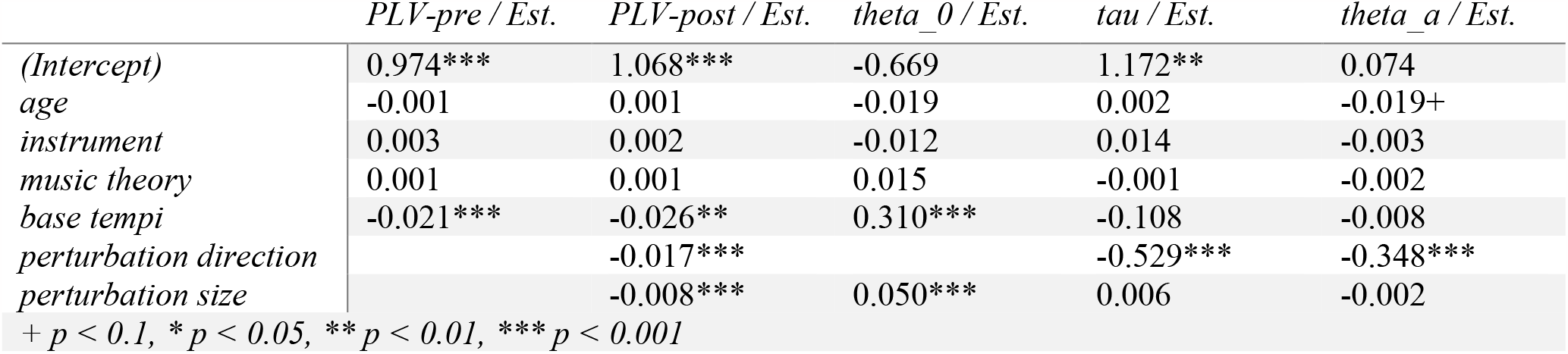
Relationship between experimental manipulation and behavioral entrainment outcomes.

|  | *PLV-pre / Est.* | *PLV-post / Est.* | *theta_0 / Est.* | *tau / Est.* | *theta_a / Est.* |
| --- | --- | --- | --- | --- | --- |
| (Intercept) | 0.974*** | 1.068*** | -0.669 | 1.172** | 0.074 |
| age | -0.001 | 0.001 | -0.019 | 0.002 | -0.019+ |
| instrument | 0.003 | 0.002 | -0.012 | 0.014 | -0.003 |
| music theory | 0.001 | 0.001 | 0.015 | -0.001 | -0.002 |
| base tempi | -0.021*** | -0.026** | 0.310*** | -0.108 | -0.008 |
| perturbation direction |  | -0.017*** |  | -0.529*** | -0.348*** |
| perturbation size |  | -0.008*** | 0.050*** | 0.006 | -0.002 |

+ $p < 0.1$ , \* $p < 0.05$ , \*\* $p < 0.01$ , \*\*\* $p < 0.001$

### Relationship between behavioral entrainment outcomes and language measurements

Based on the model outcomes (table 1), base tempi were used to model the relationship between PLV-pre and language measurements. Base tempi, perturbation direction, and perturbation size were used to model the relationship between PLV-post and language measurements. Base tempi and perturbation size were used to model the relationship between *θ*_0_ and language measurements. For *τ* and *θ*_*a*_, perturbation direction was used to model the relationship between PLV-post and language measurements.

### Relationship between phase locking values and language measurements

The full model that tested the relationship between PLV and language assessments was as follows:

~~~
model.full <- lmer(PLV-pre/PLV-post ∼ age + instrument + theory + base tempi/base tempi * perturbation size * perturbation direction * (WJ Letter-Word ID + WJ Word Attack + TOWRE Sight Word + TOWRE Phonemic Decoding + GORT Fluency + GORT Comprehension) + (1|sub), data = data)
step(model.full)
model.final = lmer(model found by stepwise regression method, data = data)
anova(model)
~~~

When PLV-pre was the model outcome, the stepwise regression method showed a significant main effect of base tempi (*b* = -0.019, *SE* = 0.006, t = -3.157, p < 0.01). There was no significant relationship between PLV-pre and language measurements.

When PLV-post was the model outcome, there was a significant main effect of perturbation size (*b* = -0.027, *SE* = 0.012, t = -2.275, p < 0.05). Interaction effects were found between perturbation size and direction (*b* = 0.038, *SE* = 0.016, t = 2.314, p < 0.05), and between perturbation direction and WJ Letter Word ID (*b* = 0.006, *SE* = 0.002, t = -2.275, p < 0.05, Figure 2a). There was also a three-way interaction of base tempi, perturbation size, and perturbation direction (*b* = -0.018, *SE* = 0.007, t = -2.494, p < 0.05), and a three-way interaction of base tempi, perturbation direction, and GORT comprehension (*b* = 0.005, *SE* = 0.002, t = 2.010, p < 0.05, Figure 2b).

**Fig. 2.**
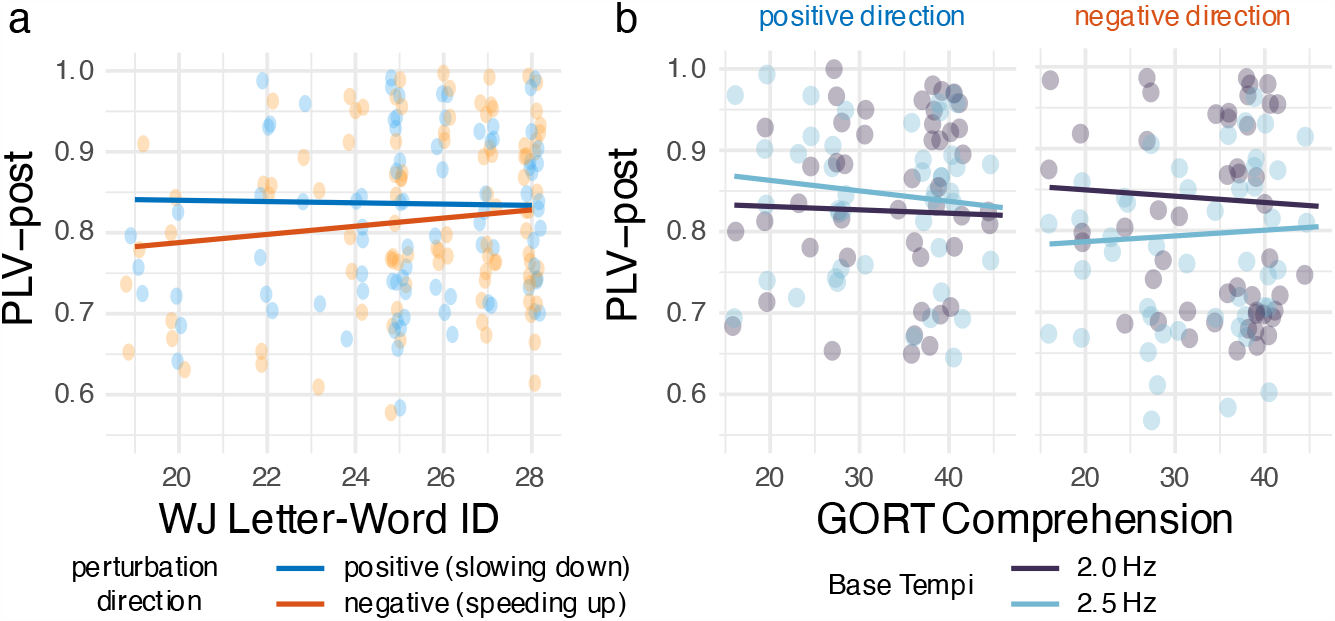
Predicted value of PLV-pre and PLV-post, for languages measures showing a significant effect.

### Relationship between curve fitting parameters and language measurements

The full model that tested the relationship between *θ*_0_ and language assessments was as follows, using normalized *SE* (*θ*_0_) as prior weights. As mentioned before, data used to model the relationship between *θ*_0_ and language assessments only included the trials when new tempi were faster than base tempi, due to the fact that participants would still tap based on the base tempo at the perturbation site when new tempi were slower than base tempi. Therefore *θ*_0_ cannot reflect the degree of behavioral perturbation for the trials that slowed down in tempo.

~~~
model.full <- lmer(*θ*_0_ ∼ age + instrument + theory + base tempi * perturbation direction * (WJ Letter-Word ID + WJ Word Attack + TOWRE Sight Word + TOWRE Phonemic Decoding + GORT Fluency + GORT Comprehension) + (1|sub), weights = normalized *SE* (*θ*_0_), data = new tempi > base tempi trials)
step(model.full) model = lmer(model found by stepwise regression method, data = data)
anova(model)
~~~

When *θ*_0_ was the model outcome, there was a significant main effect of base tempi (*b* = 7.828, *SE* = 2.341, t = 3.343, p<0.01), GORT Comprehension (*b* = 0.297, *SE* = 0.142, t = 2.094, p<0.05, fig. 3g), Interaction effects were found between base tempi * WJ Word Attack (*b* = -0.065, *SE* = 0.031, t = -2.071, p<0.05, fig. 3a), base tempi * TOWRE Sight Word (*b* = -1.068, *SE* = 0.329, t = -3.250, p<0.01, fig. 3b), base tempi * GORT Fluency (*b* = 0.014, *SE* = 0.007, t = 2.059, p<0.05, fig. 3c), base tempi * GORT Comprehension (*b* = -0.151, *SE* = 0.062, t = -2.425, p<0.05, fig. 3d), perturbation size * WJ Letter Word ID (*b* = 0.005, *SE* = 0.002, t = 0.040, p<0.05, fig. 3e), and perturbation size * GORT Comprehension (*b* = -0.013, *SE* = 0.006, t = 0.037, p<0.05, fig. 3f). Three-way interactions were found between base tempi * perturbation size * GORT Comprehension (*b* = 0.006, *SE* = 0.003, t = 2.164, p<0.05).

**Fig. 3.**
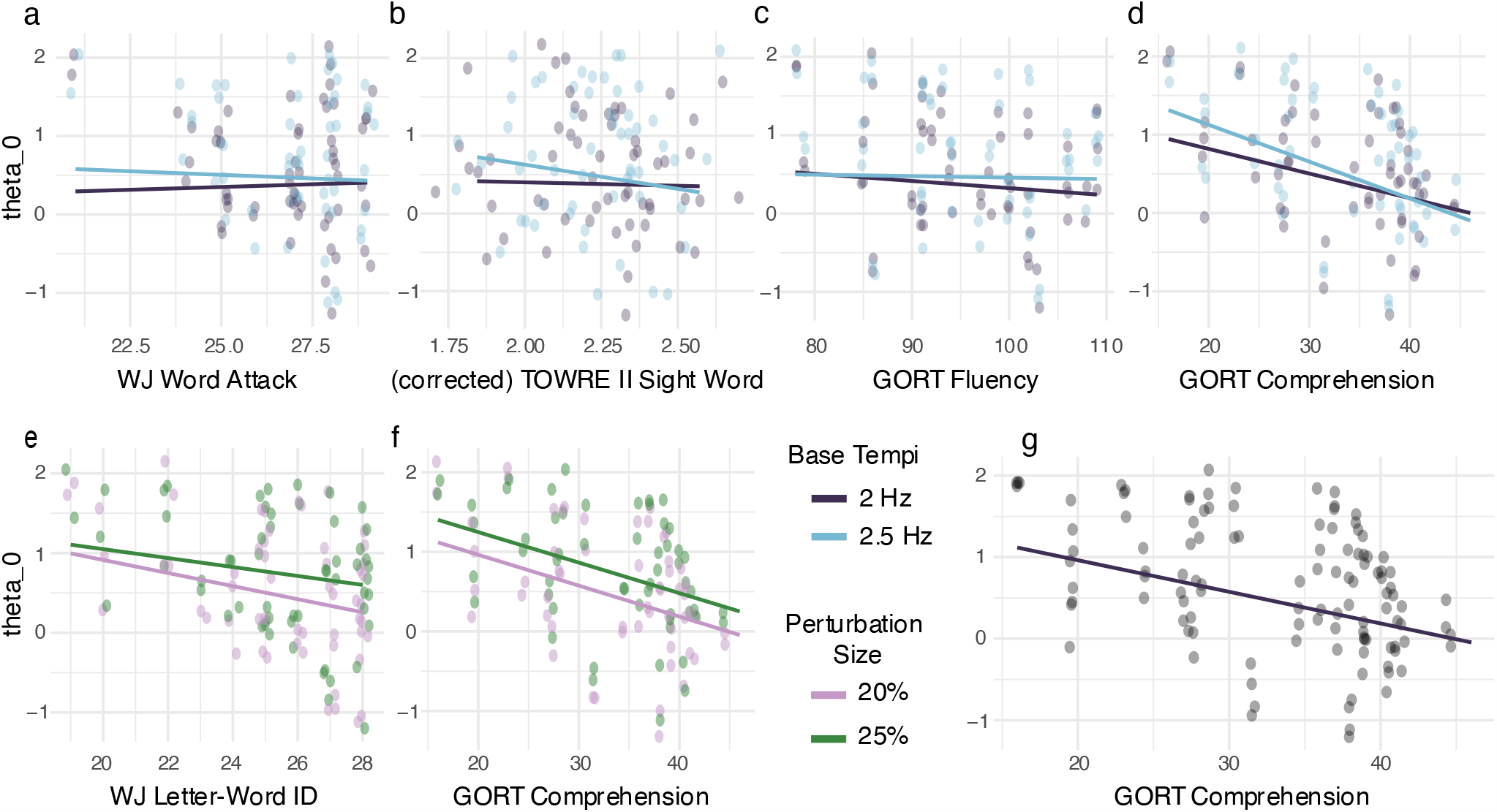
Predicted values of *θ*_*a*_, for language measurements showing a significant relationship

The full model that tested the relationship between, *θ*_*a*_ and language assessments was as follow, using normalized *SE* (*τ*/*SE* (*θ*_*a*_) as prior weights.

~~~
model.full <- lmer(*τ/ θ*_*a*_ ∼ age + instrument + theory + perturbation direction * (WJ Letter-Word ID + WJ Word Attack + TOWRE Sight Word + TOWRE Phonemic Decoding + GORT Fluency + GORT Comprehension) + (1|sub), weights = normalized *SE* (*τ*/*SE* (*θ*_*a*_), data = data)
step(model.full) model = lmer(model found by stepwise regression method, data = data)
anova(model)
~~~

When *τ* was the model outcome, significant interaction effects were found between perturbation direction and WJ Letter-Word ID (*b* = -0.067, *SE* = 0.026, t = -2.605, p<0.01, fig. 4a), perturbation direction and WJ Word Attack (*b* = 0.072, *SE* = 0.030, t = 2.428, p<0.05, fig. 4b), and perturbation direction and GORT comprehension (*b* = 0.014, *SE* = 0.006, t = 2.338, p<0.05, fig. 4c)

**Fig. 4.**
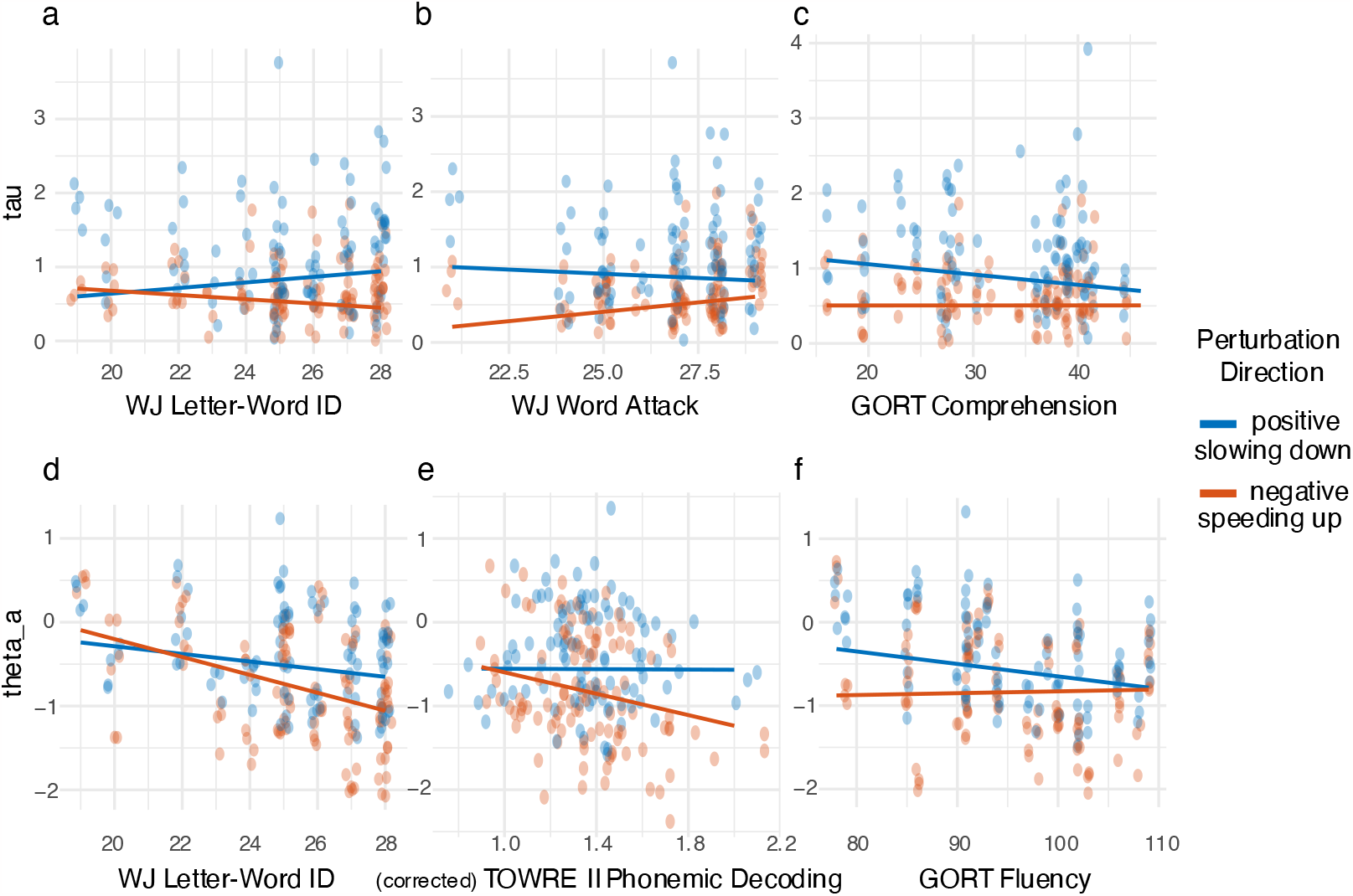
Predicted values of *θ*_*a*_, for language measurements showing a significant relationship

When *θ*_*a*_ was the model outcome, there was a significant main effect of perturbation direction (*b* = 0.563, *SE* = 0.269, t = 2.090, p < 0.05). Significant interaction effects were found between perturbation direction and WJ Letter-Word ID (*b* = -0.061, *SE* = 0.011, t = -5.541, p < 0.001, fig. 4d), perturbation direction and TOWRE Phonemic Decoding (*b* = -0.626, *SE* = 0.128, t = -4.891, p < 0.001, fig. 4e), and perturbation direction and GORT fluency (*b* = 0.017, *SE* = 0.003, t = 5.375, p < 0.001, fig. 4f). Table 2 shows the complete model outcomes of the relationship between all the behavioral analysis outcomes and reading measurements.

**Table 2.**
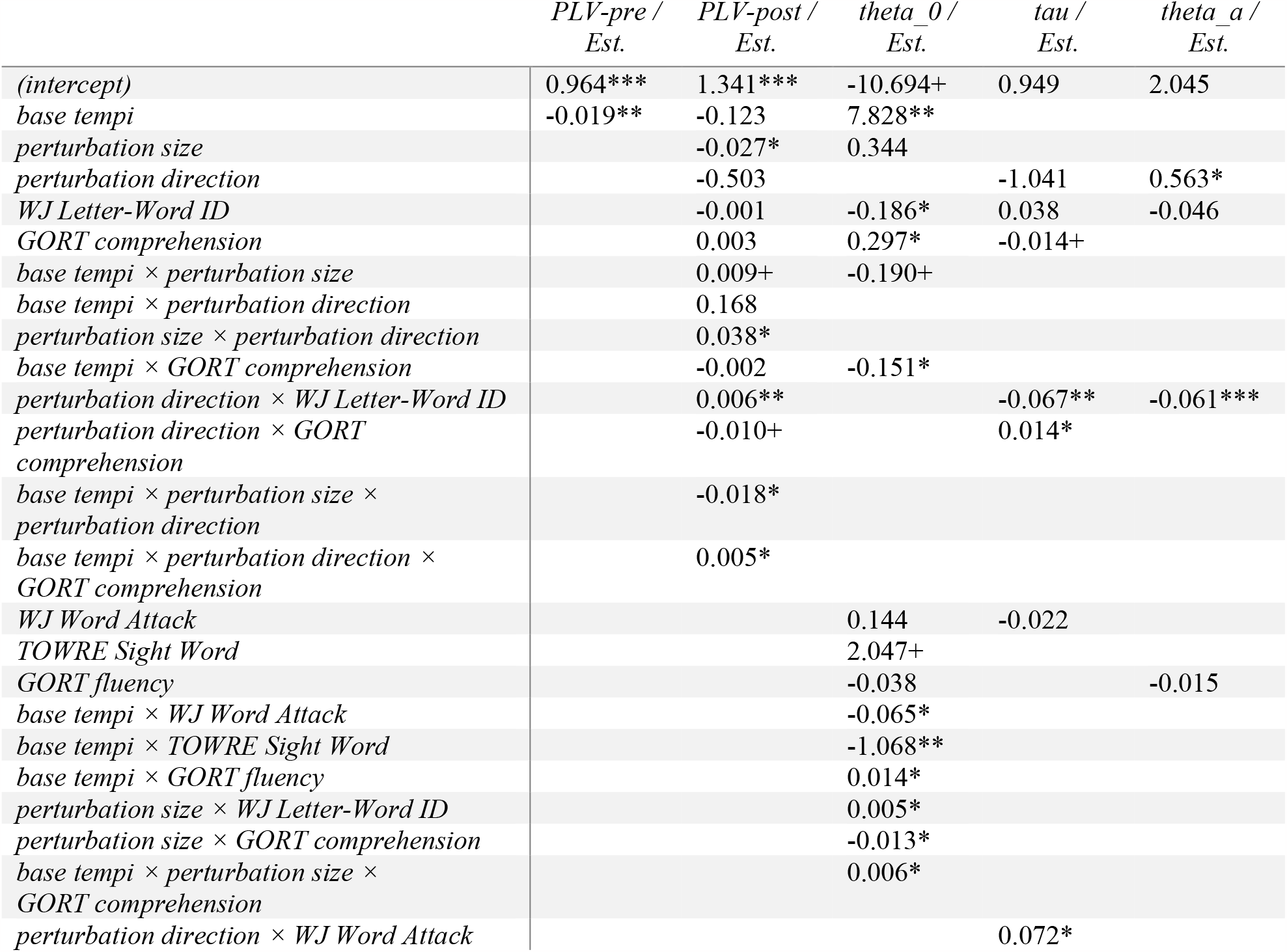
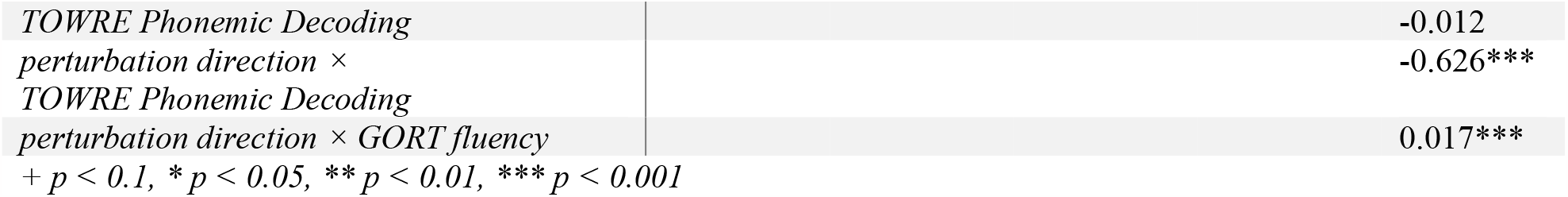
Relationship between curve fitting parameters and language measurements.

## Discussion

The current study tested the relationship between behavioral entrainment to tempo perturbation stimuli and reading measurements in 32 neurotypical adults. The first aim of this study is to address the limitation of using isochronous stimuli in past literature when study the relationship between auditory temporal processing and reading. The second aim is to apply a novel curve-fitting method to capture the nonlinearity of participants’ sensorimotor synchronization performance in response to the perturbation. This approach can provide a deeper understanding of the relationship between the dynamics of auditory temporal processing and reading measurements.

We first examined the phase locking value (PLV) during the pre- and post-perturbation time windows separately. Our regression model did not identify any relationship between participants’ phase consistency and reading measurements during the pre-perturbation time window. Several previous studies have utilized tapping to a metronome as one of the auditory processing measurements (Bégel et al., 2022; Corriveau & Goswami, 2009; David et al., 2007; Kertész & Honbolygó, 2021; Lundetræ & Thomson, 2018). Among these studies, tapping performance was indexed using either inter-tap-interval variabilities or PLV. Generally, poorer reading measurements were associated with higher inter-tap-interval variabilities or lower PLV. For instance, both Bégel et al. (2022) and Kertész & Honbolygó (2021) have found significant group differences in PLV between children with and without developmental dyslexia. The inconsistency between our findings and past research could be attributed to age differences or the fact that neurotypical adults do not exhibit sufficient individual differences in these measurements. This further suggests that measuring phase consistency using isochronous stimuli may not be sensitive enough to probe individual differences in the neurotypical adult population.

The phase locking value (PLV) during the post-perturbation time window was positively associated with the WJ Letter-Word ID task when the perturbation direction was negative. We also observed a three-way interaction concerning the relationship between PLV-post and GORT comprehension: A generally negative relationship between PLV-post and GORT comprehension, except when the base tempo is 2.5Hz and the perturbation direction was negative (speeding up). These mixed results might be attributed to a limitation of how the PLV-post was calculated, which was based on 8 taps after the perturbation onset. Participants who were more perturbed at the perturbation site but recovered faster could yield the same PLV as participants who were less perturbed but recovered slower. According to behavioral entrainment analysis, participants’ adaptation following the perturbation onset takes a roughly exponential time course (Figure 1). This is expected because tempo change engages both period correction as well as phase correction, and the exponential time course is consistent with observations in the previous tapping literature (Repp, 2005). To gain a better understanding of the adaptation from perturbation, an exponential decay model was fit to participants’ tapping relative phase during the post-perturbation time window.

Our curve fitting analysis provided more insights into the relationship between behavioral entrainment during the post-perturbation time window and reading measurements. Regression analysis on the curve fitting parameter *θ*_0_ showed that smaller phase error at the perturbed tone (smaller *θ*_0_) was associated with better scores on almost all the reading measurements, except for TOWRE phonemic decoding. Note that *θ*_0_ is not absolute individual phase error at the perturbation site; rather it reflects how much a participant’s phase is being perturbed at the perturbation site. Therefore, smaller phase error *θ*_0_ at the perturbation site indicates faster tapping adjustment to the tone that came in early unexpectedly, which indicates better sensitivity to temporal change at a fast time scale (67∼100ms in the current study). The overall negative association between *θ*_0_ and reading measurements confirms that individual differences in auditory temporal processing at a fast time scale is strongly linked to their reading measurements, in both word and pseudoword reading, and reading fluency and comprehension. The relationship between accuracy at the perturbation site and reading measurements is supported by both the Precise Auditory Timing Hypothesis framework (PATH; (Tierney & Kraus, 2014) and the OPERA hypothesis (Patel, 2011, 2014). Both frameworks emphasize the significance of precision and accuracy in processing timing information. Such precision can manifest in improved reading skills by aligning the perception of timing with meaningful phonological categories, word boundaries, and phrase boundaries.

The adaptation rate *τ* was in general positively associated with word level reading measurements (WJ Letter-Word ID and Word Attack), and negatively associated with sentence level reading measurement (GORT comprehension). Since adaptation rate *τ* reflects how fast participants recover from the perturbation site, this result might reflect different processing mechanisms at word and sentence level.

Relaxation timing accuracy *θ*_*a*_ is negatively associated with WJ Letter-Word ID, TOWRE phonemic decoding, and GORT fluency. Since *θ*_*a*_ can be considered as a measurement of negative mean asynchrony (NMA), which is an important aspect of sensorimotor synchronization, referring to the observation that taps usually precede tone onsets by a few tens of milliseconds. In the current study, participants with more negative *θ*_*a*_, which means bigger NMA (more negative, indicating tap time is more ahead of the tone onsets) had better performance on WJ Letter-Word ID, TOWRE phonemic decoding, and GORT fluency scores. Past studies have found inconsistent relationships between NMA and reading measurements. For example, Wolff (2002) found a significantly bigger NMA in children with developmental dyslexia. Leong & Goswami (2014) found bigger NMA in adults with developmental dyslexia. However, Thomson et al. (2006) found no group differences in NMA between adults with and without developmental dyslexia, while Pagliarini et al. (2020) found the opposite relationship, in which children and adults with developmental dyslexia fail to show timing anticipation compared to their control group. The fact that our study found an association between larger NMA and better performance on reading measurements in the neurotypical adult population might reflect the different nature of the populations we tested. However, the mechanism of NMA is still not entire clear. Some consider it as an anticipatory mechanism that generates predictions based on prior knowledge, and some consider it as a “time-delayed” feedback system among neural oscillators, in which case prediction is not required (Large et al., 2023; Palmer & Demos, 2022). Therefore, further experimental and theoretical work is needed to explain the mechanism behind the relationship between NMA and reading measurements.

## General discussion

A strong correlation between auditory temporal processing and reading proficiency has been consistently observed across clinical and nonclinical populations, spanning various age groups and languages. This association has spurred theoretical frameworks suggesting potential cascading effects of auditory temporal processing on the subsequent development of literacy skills. Specifically, rhythm sensitivity in both the music and speech domains is seen as a fundamental skill for accurately tracking the hierarchical acoustic components in speech. These, in turn, lay the groundwork for the development of reading skills. However, the empirical validation of this hypothesis has primarily utilized stimuli with an isochronous underlying beat structure (i.e., stationary signals). This choice of stimuli has raised concerns regarding its applicability in reflecting rhythm sensitivity in both the music and speech domains (Ozernov-Palchik & Patel, 2018).

In this context, we argue that the concern regarding the use of isochronous stationary signals does not negate the value of studying sensitivity to speech rhythm from the perspective of the music domain. Existing research has demonstrated the existence of a shared neural network for speech and music perception, justifying the use of musical stimuli to investigate rhythm sensitivity in both domains (Patel, 2011, 2014; Tierney & Kraus, 2014). However, our primary concern with isochronous stationary signals is their inability to capture the observed nonlinearity present in both speech and music perception domains. Moreover, these signals, as employed in previous studies, offer a single steady timescale for investigation, making it challenging to examine atypical auditory temporal processing occurring at faster timescales, which is also crucial in models like the temporal sampling framework.

The current study highlighted that sensorimotor synchronization during the post-perturbation time window yields significantly superior predictive value across a wide array of reading measurements when compared to sensorimotor synchronization during the pre-perturbation time window, which did not predict reading measurements. Our curve fitting analysis effectively captured the nonlinearity in participants’ tapping performance, offering additional insights into auditory temporal processing abilities in response to timing changes in auditory signals—a phenomenon that occurs frequently in speech and music signals. Overall, our observations addressed the limitations of employing a slow tempo metronome to probe the relationship between auditory temporal processing ability and reading measurements present in previous literature. This is important for two primary reasons: firstly, it allows for higher sensitivity in the neurotypical population to probe individual differences in auditory temporal processing using the sensorimotor synchronization task, enabling the exploration of the relationship between auditory temporal processing and literacy skills outside of developmental and clinical populations. Secondly, despite theories such as the Temporal Sampling Framework (TSF) proposing that atypical neural entrainment to speech rhythm may potentially disrupt auditory processing at higher frequencies, such as rise time/phonemes, through the hierarchical structure of neuronal activities, utilizing the sensorimotor synchronization task with a stationary signal at a slower frequency does not effectively allow us to simultaneously observe individual differences in auditory temporal processing across both slow and fast timescales. Nevertheless, we were able to characterize the relationship in both timescales using a non-stationary auditory stimulus design.

We acknowledge several limitations in the present study. Firstly, the application of the sensorimotor synchronization task inevitably introduces the potential confounding factor of the involvement of the motor system. One could argue that participants showing less perturbation (smaller *θ*_0_) might reflect more efficient auditory-motor system integration, rather than necessarily indicating poorer auditory temporal processing ability. We aim to address this limitation through the analysis of the neural measurements recorded in the current study. Secondly, one of the reading measurements – TOWRE – presented a ceiling effect in our population. This suggests that a different reading measurement is needed to effectively probe neurotypical adults’ individual differences in timed word-level processing ability. Thirdly, we recognize that adopting a non-stationary signal design, such as a perturbation rhythm, does not imply an assumption that the underlying mechanism is identical to processing speech rhythm—an aspect emphasized as a critical foundation for successful literacy development in TSF. Therefore the current study cannot fully account for the neural mechanism behind the relationship between auditory temporal processing and reading development. However, the significance of designing non-stationary stimuli without linguistic information lies in its capacity to characterize lower-level auditory processing in a more dynamic manner. This approach enables the observation of neural systems responding to expected and unexpected timing changes, aligning more closely with the dynamics of speech and music perception.

## Notes

**Statements and Declarations** **Competing Interests**: Authors declare no competing interests.

### Competing Interest Statement

The authors have declared no competing interest.

